# Structural Variants at Intronic CTCF Loop Anchors Drive Differential Exon Usage

**DOI:** 10.1101/2025.07.29.667294

**Authors:** Nicholas Moskwa, Minji Kim, Sabriya A. Syed, Charles Lee

**Affiliations:** The Jackson Laboratory for Genomic Medicine, Farmington, CT; Department of Computational Medicine and Bioinformatics, University of Michigan, Ann Arbor, MI

**Keywords:** Structural Variants, CTCF, chromatin loops, alternate splicing, mouse strains

## Abstract

CTCF-mediated chromatin loops are known to influence gene regulation, yet their role in pre-mRNA splicing remains incompletely understood. Here, we demonstrate that structural variants (SVs) at the anchors of intronic CTCF loops can modulate exon usage. By integrating high-resolution three-dimensional (3D) genome organization and gene expression datasets from C57BL/6J (B6) and 129S1/SvImJ (129S) mouse embryonic stem cells (ESCs), with structural variant (SV) maps from the 129S mouse, we identified thousands of intron-anchored CTCF loops. Our data indicate that SVs intersecting loop anchors are more frequently associated with differential exon inclusion events than with changes in overall gene expression. CRISPR/Cas9 deletion of two SV-harboring intronic CTCF sites in *Numbl* and *Ireb2* validated the predicted splicing shifts observed between B6 and 129S ESC that correspond with diminished long-range chromatin looping. Our findings reveal a direct mechanistic link between 3D genome architecture and alternative splicing and highlight non-coding SVs as modulators of transcript diversity. Our study has thus identified a novel class of CTCF-bound regulatory elements regulating alternative splicing. Cataloging and validating these functional elements will elucidate molecular mechanisms underlying phenotypic variation within populations.

## INTRODUCTION

The three-dimensional (3D) architecture of the genome is fundamental to gene regulation and cellular identity. Chromatin folding shapes the transcriptional landscape by organizing DNA into hierarchical structures that control the physical proximity of regulatory elements (Hsieh et al., 2020; Mohana et al., 2023). At the megabase scale, chromatin is partitioned into topologically associating domains (TADs), which constrain enhancer-promoter interactions within domains and insulate them from neighboring regions (Kraft et al., 2019; Lupiáñez et al., 2015). Within TADs, chromatin loops bring distal regulatory elements, such as enhancers and promoters, to facilitate gene expression. These loops are orchestrated by architectural proteins including cohesin, RNA Polymerase II (RNAPII), Yin Yang 1 (YY1) and CCCTCF binding factor (CTCF). CTCF serves a dual role in both TAD boundary formation and local loop mediation (Kubo et al., 2021; Levo et al., 2022; Weintraub et al., 2017).

Large-scale enhancer mapping efforts, such as those by the ENCODE and 4D Nucleome consortia, have revealed the vast regulatory complexity of mammalian genomes (The ENCODE Project Consortium et al., 2020). Over a million enhancer-like elements have been catalogued across diverse human and mouse cell types (The ENCODE Project Consortium et al., 2020), emphasizing that genes are controlled by context-specific noncoding regulatory elements (Heinz et al., 2015). Importantly, genetic variation in these regulatory elements has been linked to individual differences in gene expression and chromatin organization (Gorkin et al., 2019; Grubert et al., 2015; Reddy et al., 2012; Tang et al., 2015; The ENCODE Project Consortium et al., 2020). Chromatin organization can be assessed with chromatin conformation capture-based methods, including Hi-C and chromatin interaction analysis with paired-end tag sequencing (ChIA-PET). Hi-C assesses the frequency of physical association between DNA fragments while ChIA-PET involves immunoprecipitating proximity-ligated chromatin fragments to map protein-mediated chromatin interactions. Integration of the enhancer catalogs and genome organization datasets have indicated that genetic variation can rewire 3D chromatin architecture and contribute to functional phenotypic differences.

CTCF is a key chromatin architectural protein known for mediating long range chromatin loops through sequence-specific DNA binding and homodimerization events (Kubo et al., 2021; Mohana et al., 2023; Tang et al., 2015). While extensively studied for its roles in organizing TADs and facilitating promoter-enhancer interactions, a substantial fraction (22%-29%) of CTCF binding sites actually occur within introns (Kim et al., 2007). However, these intronic CTCF sites remain poorly characterized. Some studies have suggested that they may play a role in alternative splicing regulation. For instance, RNA splicing quantitative trait loci (QTL) often overlap with CTCF motifs (Li et al., 2016), while CTCF haploinsufficiency in mice has been associated with widespread intron retention (Alharbi et al., 2021). Additionally, CTCF binding at exonic regions has been associated with transcriptional pausing and exon inclusion (Shukla et al., 2011) and Ruiz-Velasco et al. (2017) correlated intragenic CTCF looping with exon inclusion events. However, to date, the functional consequence of intronic CTCF loops has not been tested directly *in vitro* or *in vivo*.

Here, we directly test the functional role of intronic CTCF sites in alternative splicing. We leverage natural genetic variation between two common mouse strains (C57BL/6J (B6) and 129S1/SvImJ (129S)) to identify intronic CTCF-bound chromatin loops that are disrupted by a structural variant (SV) in one strain and not the other. A subset of these SV-disrupted loops is associated with strain-specific exon usage. To validate this, we used CRISPR/Cas9 to delete SVs overlapping intronic CTCF-bound elements and showed that loss of CTCF occupancy leads to changes in exon inclusion. These findings provide evidence that intronic CTCF-mediated loops regulate alternative splicing.

## RESULTS

### Chromatin looping differs significantly between B6 and 129S ESC

To investigate whether genetic variation at CTCF-bound loop anchors contributes to differences in gene expression, we utilized publicly available in-situ CTCF ChIA-PET data from ENCODE (Wang et al., 2021; The ENCODE Project Consortium et al., 2020; Dunham et al., 2012). In-situ CTCF ChIA-PET data sequenced at 300 million (M) reads per sample identified 439,938 CTCF peaks in 129S and 322,914 peaks in B6 ESCs (approximately sevenfold more than previous ChIP-seq studies (e.g., (Nora et al., 2017)), possibly due to the ten-fold increase in sequencing read depth (300M reads vs 30M reads)). ChIA-PET loop calling identified 230,685 loops in 129S and 232,159 loops in B6 ESCs. To determine strain specificity, we used BEDTools (Quinlan & Hall, 2010) to compute the symmetric difference between loop sets, identifying 201,449 129-specific loops and 203,796 B6-specific loops (**Figure 1A**). A complementary analysis with diffloop (Lareau & Aryee, 2018) yielded similar results: 198,407 129-specific loops and 190,874 B6-specific loops (padj ≤ 0.1, |log₂FC| ≥ 0.5), with substantial overlap between methods (93% diffloop overlap with the 129-specific loops and 95.6% diffloop overlap with the B6-specific loops). We next filtered for loops with significant CTCF binding at the loop anchors (q-value < 0.05), yielding 158,890 CTCF-bound loops (CBLoops) in 129S and 192,277 in B6 (**Figure 1A**). Notably, 80% of B6-specific and 41% of 129S-specific CBLoop anchors contained a canonical CTCF motif.

**Figure 1.**
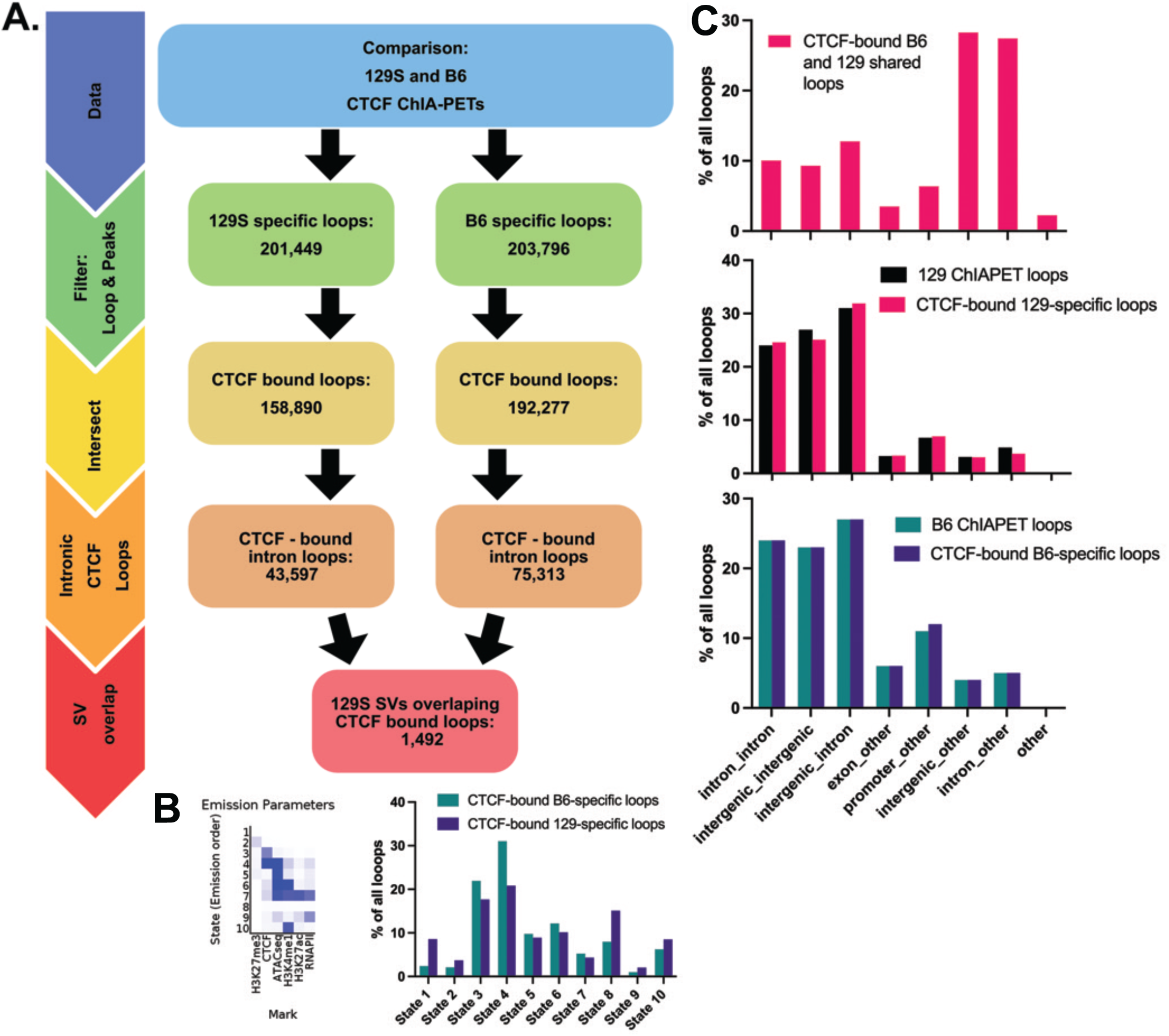
Data analysis schema for prioritizing SVs potentially impacting genome organization. **(A)** The workflow selects for chromatin loops unique to each mouse strain. Loops are then stratified by selecting for those with significant CTCF peaks. Those are then overlapped with SVs identified from a B6 and 129S comparison; **(B)** ChromHMM analysis identifies 10 distinct emission states characterized by unique histone modification presence/absence. The bar plot to the right displays the % of loops with each emission state characteristics. **(C)** Bar plots summarizing loop anchor annotation for CTCF-bound 129-specific, B6-specific and 129 and B6 shared loop lists.

### CBLoops are regulatory elements with features distinct from enhancers and TAD boundaries

To determine whether CBLoops overlap known cis-regulatory elements, we performed ChIPseq for histone modifications H3K27ac, H3K4me1, H3K27me3, and H3K4me3 in both strains. ChromHMM analysis that included ChIPseq datasets and open chromatin (ATAC-seq) regions (Ferraj et al., 2023) found that ∼50% of strain-specific CBLoops lacked clear histone modification signatures, while ∼20% exhibited a “poised enhancer” profile (H3K4me1-positive, H3K27ac negative observed in emission states 6 and 7) (Creyghton et al., 2010) (**Figure 1B**). Only 1.2% (129S) and 2.5% (B6) of CBLoop anchors coincided with TAD boundaries, as defined by diamond insulation scores using cooltools (Open2C et al., 2024). These data suggest that CBLoop form between uncharacterized noncoding regions, distinct from typical promoter-enhancer loops or TAD boundaries (Kubo et al., 2021; Levo et al., 2022; Weintraub et al., 2017).

### Intronic CBLoop anchors commonly contain B2 SINE elements and a subset is prone to structural variation

A substantial proportion of CBLoops were intronic: 43,597 in 129S (27% of CBLoops) and 75,313 in B6-specific (39% of CBLoops) (**Figure 1C**). ChIA-PET tag density plots visualizing CTCF binding at these intronic sites confirmed high CTCF occupancy at the CBLoop anchors (**Supplemental Figure 1A-B** showing the intron-intron subset). HOMER motif analysis revealed strong enrichment for CTCF-related motifs (including the CTCF-like protein BORIS (Pugacheva et al., 2015), and additional strain-specific transcription factors, such as MyoD in 129S (21.6% of anchors) and ZNF341 in B6 (26.8%) (**Supplemental Figure 1C**).

We next asked whether SVs could account for CBLoop differences. Of 59,352 known SVs distinguishing the 129S mouse from the B6 reference (Ferraj et al., 2023), 1,492 overlapped intronic CBLoop anchors (**Figure 1A; Supplemental Table 1**). Motif analysis (HOMER and MEME) identified enrichment of CTCF, BORIS, and ZNF143 motifs in these SVs. Notably, 38% of the 1,492 SVs harbored the CTCF motif, with closer to 44% harboring the BORIS motif (**Supplemental Figure 2A)**. RepeatMasker analysis and findings from the CENSOR database supported these findings (Kohany et al., 2006; Smit et al., 2013) - 30% of the 1,492 SVs contained B2 SINE elements, consistent with prior studies linking B2 SINEs to CTCF loop anchors (Choudhary et al., 2023) (**Supplemental Figure 2B-C**).

Annotation against ENCODE Candidate cis-Regulatory Elements (cCREs) showed that 382 (25.6% of the SVs) of the intronic CBLoops overlapped ‘CTCF-only’ regions, 419 (28.8% of the SVs) mapped to distal enhancer-like elements, and only 190 (12.7% of the SVs) overlapped ‘promoter-like’ or ‘proximal enhancer-like’ or ‘DNase-H3K4me4’ regions. Together, these results suggest that many intronic CBLoops intersect SVs that disrupt poorly characterized cis-regulatory elements enriched for retrotransposons.

### Structural variation at CBLoop anchors is linked to alternative exon inclusion events

To assess the functional impact of CBLoop-disrupting SVs, we generated RNA-seq data from 129S (n=4) and B6 (n=3) ESCs clones (Czechanski et al., 2014). Of the 1,492 SVs, 798 overlapped genes directly, and an additional 1,217 were connected to genes via CBLoops. Differential gene expression (DEseq2) (Love et al., 2014) identified only 312 (16%) differentially expressed genes (qvalue ≤ 0.1, |log₂FC| ≥ 0.4) (**Figure 2A**, top panel). In contrast, differential exon usage (DEXseq) analysis (Anders et al., 2012) identified 9,009 differentially used exons (pvalue ≤ 0.1, |log₂FC| ≥ 0.4) existing in 1,282 (64%) genes linked to our SVs of interest via CBLoops (**Figure 2B**, top panel, **Supplemental Table 2**).

**Figure 2.**
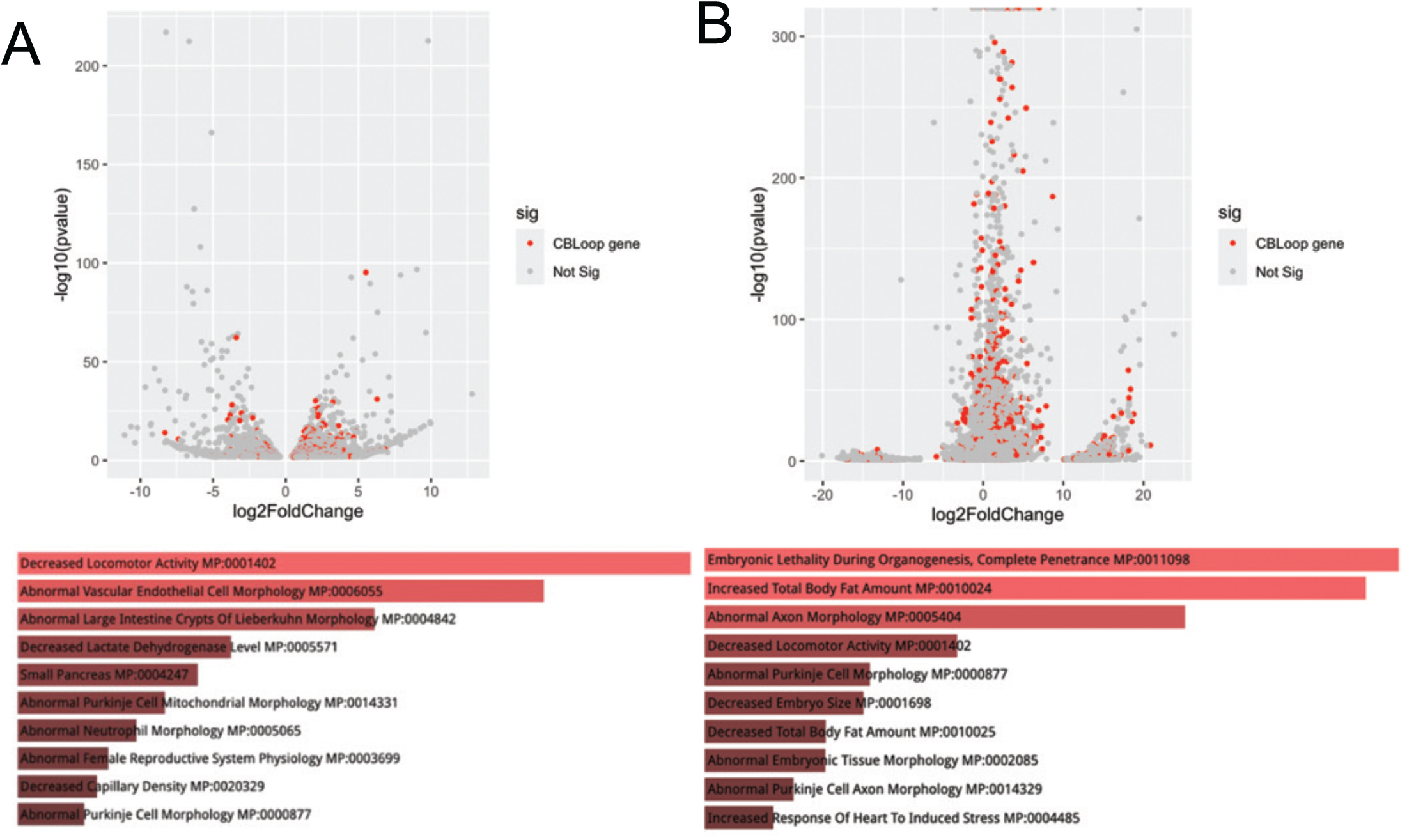
Transcriptional differences between the 129S and B6 mouse strains. **(A)** *Top*. Volcano plot displaying genes with significant differential expression (padj < 0.1, |log₂FC| ≥ 0.4). *Bottom*. Enrichr Mammalian Phenotype Ontology analysis identifies key phenotypes associated with genes impacted. **(B)** *Top*. Volcano plot displaying genes with significant differential exon usage (pvalue < 0.1, |log₂FC| ≥ 0.4). *Bottom*. Enrichr Mammalian Phenotype Ontology analysis identifies key phenotypes associated with genes impacted.

Ontology enrichment analysis (EnrichR, Mammalian Phenotype Ontology (Baldarelli et al., 2024)) revealed distinct functional categories for genes affected by differential expression versus splicing (**Figure 2A, 2B**, bottom panels), underscoring separate regulatory mechanisms.

### Intronic SVs correspond to loss of CBLoops and exon usage differences in 129S ESC

To validate candidate SVs regulating differential exon usage events, we prioritized events with strong CTCF binding differences (signal >5, –log₁₀q >150), convergent CTCF or CTCF–BORIS motifs, and associated exon usage changes (p < 0.1). One such variant, SV chr7-26971377-DEL-261, is a 261 bp deletion in the *Numbl* gene intron in 129S ESC, overlapping a B6-specific CBLoop anchor (**Figure 3A**, green highlight). ChIP-qPCR confirmed higher CTCF binding in B6 ESCs compared to 129S (**Figure 3B**), and genotyping verified the deletion in 129S clones (**Figure 3C**). ChIA-PET data revealed three CBLoops associated with the SV: an internal Numbl loop, a long-range loop connecting Numbl to Coq8b (**Figure 3A**), and an additional CBLoop linking the SV to an intron in gene Sptbn4 (**Supplemental Figure 3A**). Diffloop paired-end tag (PET) counts show loss of all three loops in 129S ESCs that have SV chr7-26971377-DEL-261 **(Figure 3D, Supplemental Figure 3B**). DEXSeq analysis showed differential exon usage at *Numbl* and *Sptbn4* exons; these genes are linked together by a CBLoop with convergent CTCF motifs at the loop anchors (**Supplemental Figure 3C–D**, yellow highlight indicates exons with pvalue < 0.1).

**Figure 3.**
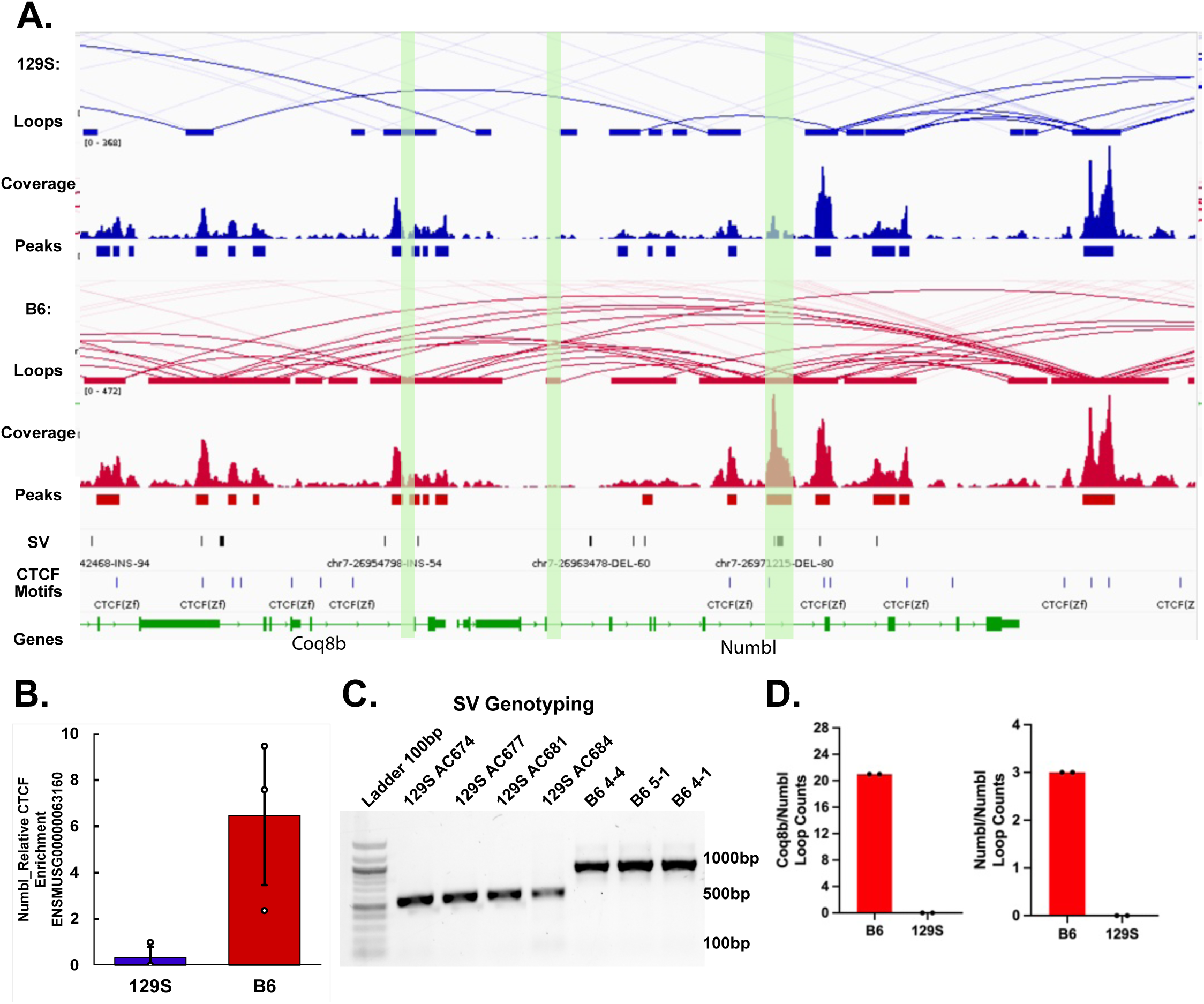
Numbl exhibits differential CTCF-bound chromatin loops between 129S and B6 mouse strains. **(A)** Genomic tracks for Numbl gene. Highlighted in green is a CTCF-bound loop anchor that is B6 specific and interrupted by an SV in the 129S ESC; **(B)** Bar plot for CTCF ChIP-qPCR confirms higher CTCF enrichment in the B6 ESCs (n=3) in comparison to the 129S ESCs (n=3); **(C)** Gel electrophoresis for genotyping PCR. B6 (n=4) and 129S (n=3) ESC clones showing a 261 bp deletion (chr7-26971377-DEL-261) in the 129S ESC clones; **(D)** Diffloop analysis shows loss of loops interacting with Numbl SV chr7-26971377-DEL-261, specifically in the 129S ESC clones.

### CRISPR/Cas9 Disruption of SVs Reproduces Splicing Effects in genes Numbl and Ireb2

To directly test the regulatory role of SV chr7-26971377-DEL-261, we introduced the deletion into B6 ESCs using CRISPR/Cas9 with Homology-Directed Repair (HDR) (Pantazis et al., 2022) (**Figure 4A**). Edited clones were confirmed by PCR and Sanger sequencing (**Figure 4B-C**), with 42% editing efficiency. Total *Numbl* expression remained unchanged (**Figure 4D**), but RNA-seq revealed altered exon usage in gene *Sptbn4* in the edited clones, specifically at an exon flanking the CBLoop anchor (**Supplemental Figure 4B**). *Coq8b and Numbl* showed no differentially used exons (**Supplemental Figure 4A**). A loss of chromatin looping between *Numbl* and *Sptnb4* was observed in the B6 and 129S ChIA-PET comparison, supporting a model in which long range CBLoops regulate differential exon usage. Together, these results indicate that the intronic CBLoop anchors in *Numbl* influence *Sptbn4* exon inclusion via chromatin architecture.

**Figure 4.**
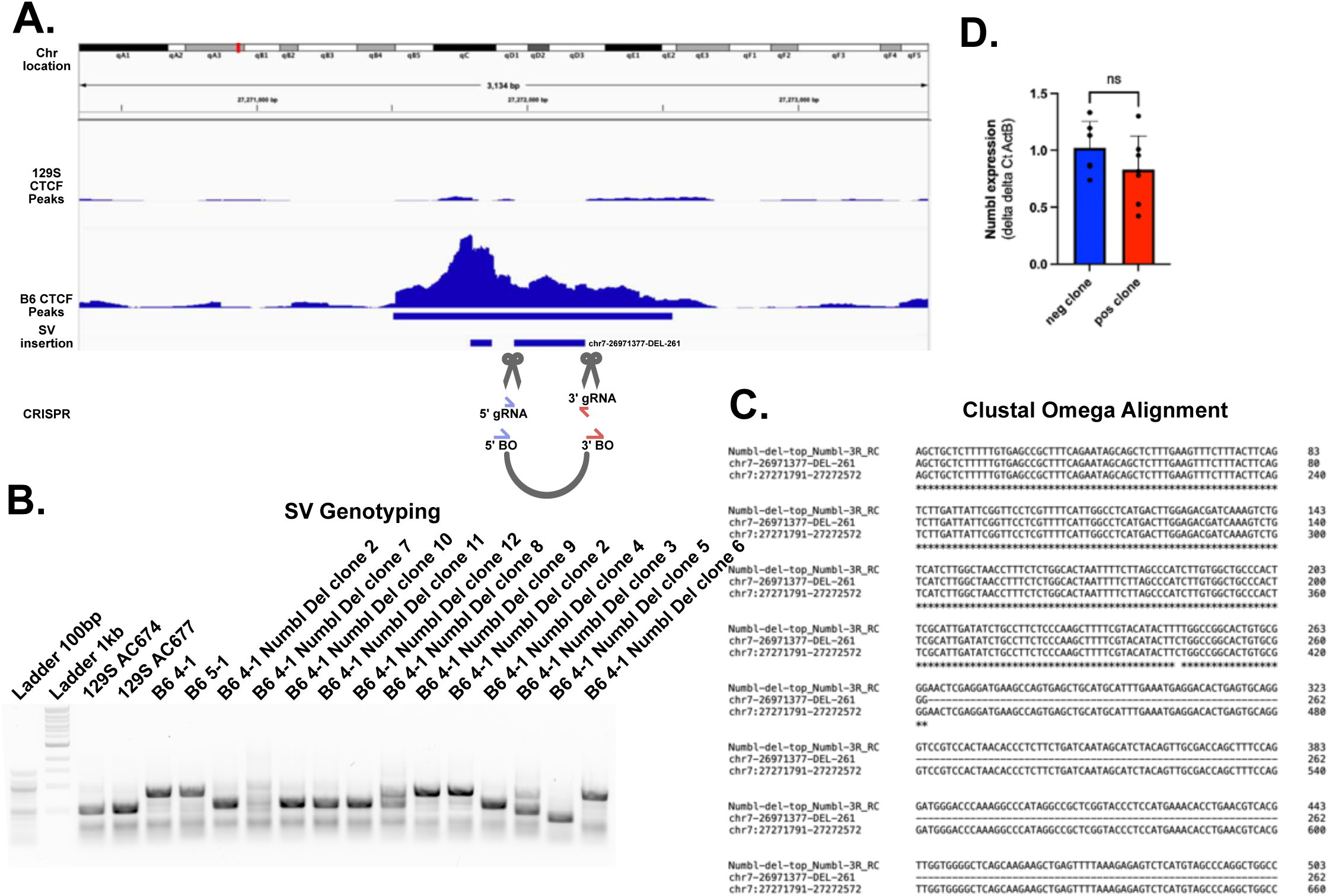
Numbl’s intronic CTCF chromatin binding site deletion in B6 ESCs using CRISPR/Cas9. **(A)** IGV schematic showing chr7:27271951-27272212 containing Numbl intron and SV chr7-26971377-DEL-261. Scissors show where on the genome CRISPR is planned to cut the DNA based on the gRNA. The grey loop shows how the bridge oligo (BO) will link the two genomic ends causing the deletion. The subsequent deletion spans > 80% of the original SV and covers the major CTCF peak; **(B)** Gel electrophoresis genotyping PCR showing unedited B6 and 129S mouse ESC compared to CRISPR edited B6 cells. The Numbl edited cells show the desired deletion band that size matches the 129S SV band; **(C)** Clustal Omega Alignments for all the Sanger sequences show the Numbl CRISPR deletion in B6, spanning 261 basepairs as expected; **(D)** RT-PCR showing no change in Numbl gene expression between unedited clones (n=6) and edited clones (n=6).

We next validated SV chr9-54812235-DEL-668 in *Ireb2*, which exists in the 129S ESC and overlaps an intronic CBLoop with strong CTCF binding and convergent motifs (**Supplemental Figure 5**). This SV is adjacent to exon 20, one of three exons showing significant usage differences (p<0.1) in the B6 and 129S ESC comparison. CRISPR/Cas9 mediated deletion of the SV in B6 ESCs was confirmed by PCR and Sanger sequencing (**Figure 5A-C**). RT-PCR demonstrated altered inclusion of exon 20 and flanking exons in the edited clones (**Figure 5D**).

**Figure 5.**
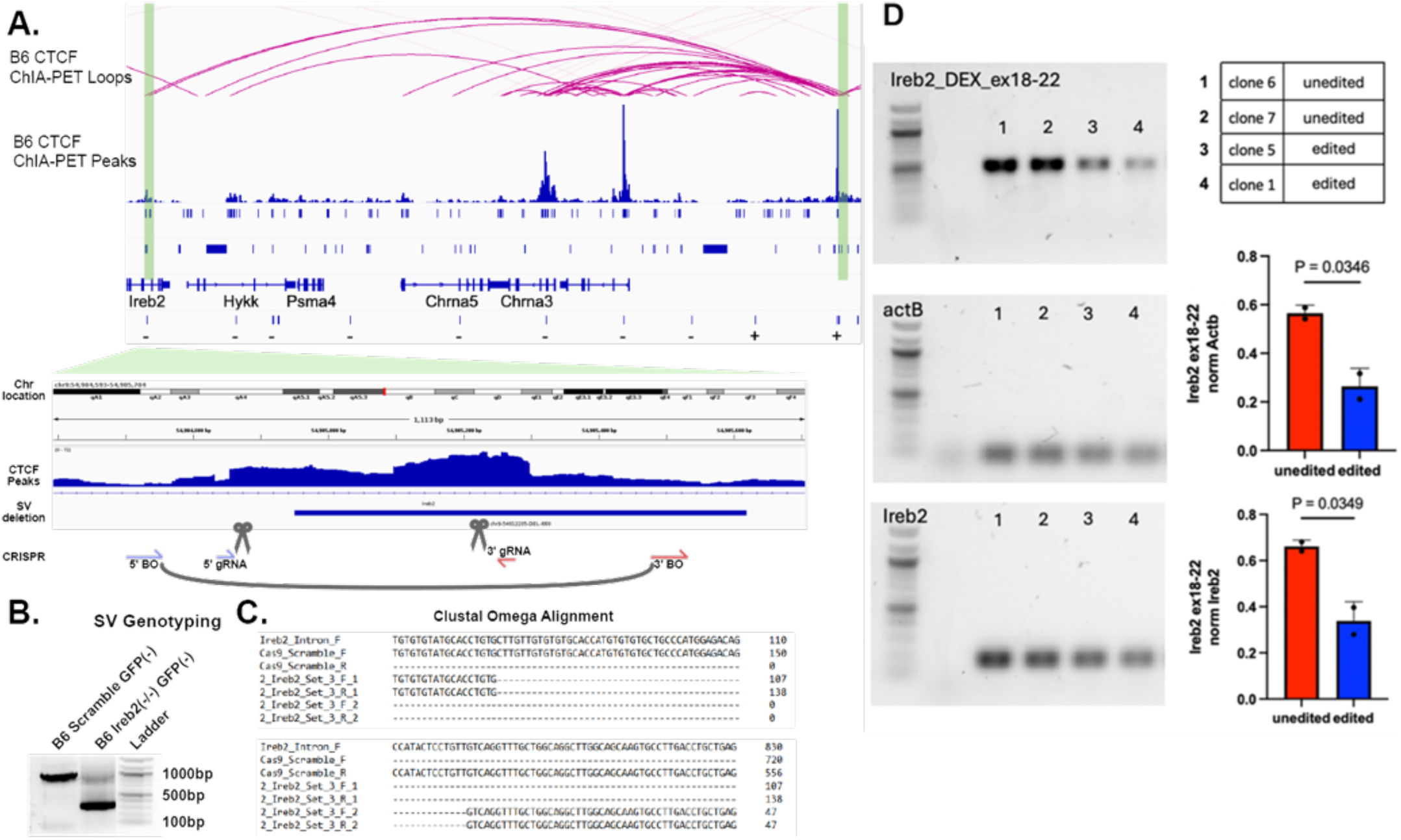
Ireb2’s intronic CTCF chromatin binding site deletion in B6 ESCs using CRISPR/Cas9. **(A)** IGV schematic showing chr9:54,898,802-54,949,866 containing SV chr9-54812235-DEL-668 in an intron of *Ireb2*. The region targeted for CRISPR/Cas9 deletion is highlighted and zoomed in. Scissors show where on the genome CRISPR is planned to cut the DNA based on the gRNA. The grey loop shows how the bridge oligo (BO) will link the two genomic ends causing a deletion in the B6 ESC. The subsequent deletion spans > X% of the original SV and covers the major CTCF peak; **(B)** Gel electrophoresis genotyping PCR showing unedited B6 and edited B6 ESC. The Ireb2 edited cells show the desired deletion that size matches the B6 unedited cells. CRISPR deleted is approximately 668 bp in size; **(C)** Clustal Omega Alignments for all the Sanger sequences show the Ireb2 CRISPR deletion; **(D)** RT-PCR shows significant difference in exons 18-22 expression when compared to ActB or a stable Ireb2 construct (exon 7-9).

Taken together, in this study, we demonstrate that SVs disrupting intronic CBLoops can modulate alternative splicing via long range chromatin looping. A proposed mechanism is displayed in **Figure 6**. CBLoops likey represent a previously underappreciated class of regulatory sequences that contribute to transcriptomic diversity and phenotypic variation between mouse strains.

**Figure 6:**
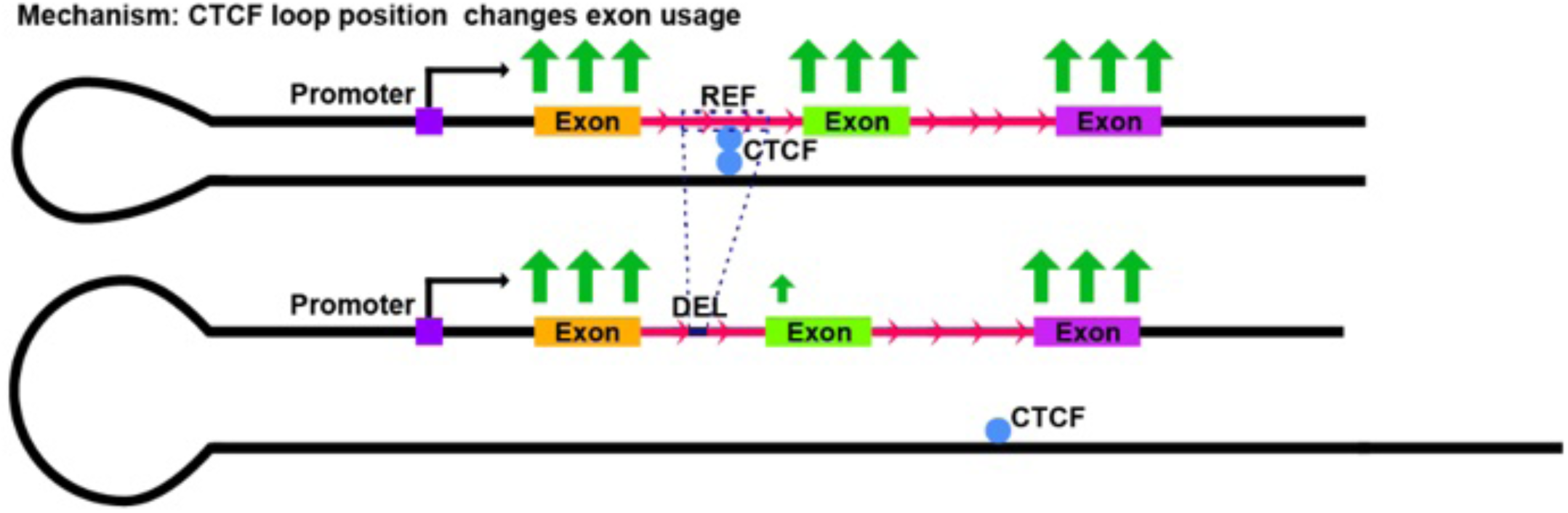
Proposed mechanism of how CTCF loop position changes exon usage. Genetic variation in gene introns can disrupt CTCF-mediated chromatin looping events. This can in turn offset exon utilization.

## DISCUSSION

CTCF is a central architect of 3D genome organization, classically known for demarcating TADs and facilitating enhancer-promoter interactions. Yet, only a minority of CTCF binding sites localize to TAD boundaries. The vast majority (especially those scattered across noncoding regions) remain functionally underexplored. Increasing evidence shows that genetic variants can alter CTCF binding and chromatin looping, thereby reshaping genome architecture and affecting gene expression. Notably, rare SVs in humans that disrupt CTCF boundaries have been linked to enhancer miswiring and developmental disorders (Lupiáñez et al., 2015), while population-level variation at CTCF binding sites has been associated with gene regulatory differences (Gorkin et al., 2019). However, whether such variants can drive phenotypic divergence between populations has remained an open question.

A substantial fraction (22-29%) of CTCF sites are located within introns. While enhancers have been identified within introns (Rigau et al., 2017), intronic regulatory elements remain poorly annotated. In this study, we systematically characterized intronic CBLoops and demonstrated their functional relevance in regulating exon inclusion. We identified strain-specific intronic loops in mouse ESCs and found that many of these loops overlapped SVs that distinguish the B6 and 129S mouse strains.

Prior studies have shown that CTCF binding within introns and exons likely modulates RNAPII elongation, thereby affecting exon inclusion (Ruiz-Velasco et al., 2017; Shukla et al., 2011). Here, we leveraged naturally occurring SVs to identify CTCF-mediated alternative splicing events that likely contribute to transcriptomic (and potentially phenotypic) divergence between the B6 and 129 mouse strains. By integrating CTCF ChIA-PET, RNA-seq, and high resolution SV maps, we identified 1,492 SVs overlapping intronic CTCF chromatin binding sites. Most of these SVs fell outside canonical enhancer regions, suggesting they disrupt novel classes of CTCF-bound regulatory elements. Moreover, many overlapped retrotransposon-derived sequences such as B2 SINEs, underscoring the complex interplay between repetitive DNA and genome regulation.

Functionally, we demonstrate that SVs disrupting intronic CBLoops (such as chr7-26971377-DEL-261 in *Numbl* and chr9-54812235-DEL-668 in *Ireb2*) can alter exon usage without necessarily affecting total transcript levels. Our findings provide direct evidence that intronic CBLoops serve as spatial regulators of alternative splicing and that SVs can perturb this regulatory mechanism. To test this, we used CRISPR/Cas9-mediated genome editing in B6 ESCs, to introduce the *Numbl* SV (chr7-26971377-DEL-261). This engineered deletion disrupted local chromatin loops and led to changes in exon inclusion in Numbl itself and the loop-connected gene *Sptbn4*.

*Numbl* is critical for cortical development. Mice lacking both *Numb* and *Numbl* exhibit embryonic lethality by E12.5, whereas *Numb* knockout mice survive (Petersen et al., 2002). These data suggest that *Numbl* is essential for viability, likely influencing critical mouse phenotypes. Therefore, understanding the effects of SVs in *Numbl*, particularly in 129S mice, could inform mouse strain selection in knockout and knockdown studies involving the nervous system. Similar caution is warranted in hepatic studies involving *Ireb2,* which harbors the SV chr9-54812235-DEL-668. *Ireb2* encodes an iron responsive element binding protein. Our results suggest that the 668 bp deletion in 129S mice affects exon inclusion through disruption of chromatin looping. This splicing change may be linked to known hepatic iron metabolism defects observed in 129S mice (Rogers, 2018).

Our study uncovers an underrecognized mechanism by which genetic variation can influence gene regulation: through disruption of chromatin loops anchored within introns. We show that intronic CTCF-mediated loops are functional regulatory structures that shape exon usage, and that SVs disrupting these loops can alter splicing patterns. This highlights a critical role for intronic regions, often overlooked in genome annotations, as scaffolds for regulatory chromatin architecture. Notably, many of the SVs we identified do not overlap known regulatory elements, yet intronic sequences in particular can harbor key cis-regulating features that modulate transcript isoform diversity.

By integrating comparative genomics, chromatin conformation data, and functional genomic approaches, we present a generalizable framework for identifying and validating noncoding regulatory elements that act through 3D genome architecture. These findings suggest that SVs and other noncoding variants affecting intronic chromatin loops may contribute to inter-individual differences in gene expression by altering splicing regulation. As such, accounting for 3D genome organization in variant interpretation could improve our understanding of the functional consequences of non-coding genetic variation in humans.

## MATERIALS AND METHODS

### ChIA-PET analysis

ChIA-PET datasets were analyzed using the ChIA-PIPE pipeline (Lee et al., 2020). Paired-end reads containing the bridge linker were extracted, and linkers were trimmed before alignment to the mm10 reference genome using BWA-MEM. Non-redundant paired-end tags (PETs) with MAPQ ≥ 30 were retained in BAM format. These bam files were processed into 2D contact maps using Juicer tools and were also used for peak and loop calling with MACS. Resulting chromatin loops and CTCF peaks were visualized using the Integrative Genomics Viewer (IGV).

### ChIA-PET TAD analysis

To assess topologically associating domains (TADs), ChIA-PIPE-generated .hic files were converted to .cool format using hic2cool. Files were balanced using Cooler’s iterative correction algorithm. TAD boundaries were identified using the cooltools diamond-insulation function (Abdennur & Mirny, 2020; Hsieh et al., 2020; Krietenstein et al., 2020).

### Loop differential analysis

Loop level differential analysis was performed using the *diffloop* R package (Lareau & Aryee, 2018).

### ESC cell culture

ESC lines derived from individual blastocysts (Skelly et al., 2020) were maintained on gelatin-coated plates (STEMCELL 7903) in ESC media composed of Dulbecco’s Modified Eagle Medium (DMEM) supplemented with 15% fetal bovine serum (FBS), 100 U/mL Penicillin-Streptomycin, 2 mM GlutaMAX, 0.1 mM non-essential amino acids (ThermoFisher Scientific 11140050), 1 mM sodium pyruvate (Fisher Scientific 11360070), 0.1 mM 2-mercaptoethanol, 500 pM LIF (R&D Systems 8878-LF-025/CF), 1 uM PD0325901 (STEMCELL 72184), and 3 uM CHIR99021 (Millipore SML1046-5MG)). Cells were grown to 70-90% confluency and passaged with 0.05% Trypin-EDTA (Fisher Scientific 25300052). During passaging, cells were counted using a hemocytometer and separated into 10 million cell aliquots for fixation or RNA extraction.

### SV genotyping PCR

SV-flanking primers were designed using NCBI Primer-BLAST. Genomic DNA was isolated using the DNeasy Blood and Tissue kit (QIAgen 69504). PCR was performed with Quick-Load® Taq 2X Master Mix (New England Biolabs M0271L) and resolved via 1% agarose gel (Invitrogen 16500500) supplemented with SYBRsafe (Invitrogen S33102). SV genotyping primer sequences can be found in **Methods, Table 1**.

### RNA extraction and expression analysis

Total RNA was extracted using the RNeasy Mini Kit (Qiagen 74104) and stored at −80°C. cDNA synthesis was performed using the QuantiTect Reverse Transcription Kit (QIAGEN 205313). Expression qPCR was performed with PowerUp SYBR Green Master Mix (ThermoFisher Scientific A25777), after samples were diluted to 10ng/ul and mixed with gene expression primers at 200 nm concentration, using a C1000 Touch™ Thermal Cycler (Bio-Rad). Relative gene expression was calculated using the 2(−ΔΔC(T)) method.

### CRISPR/Cas9 targeted deletion via nucleofection

The CRISPR/Cas9 protocol was adapted from a human iPSC protocol (Skarnes et al., 2019). sgRNAs (Synthego) and single strand oligonucleotides (ssODN; IDT) were designed using JAX CRISPR designer (https://wge.jax.org/) and resuspended overnight at RT in TE buffer at concentrations of 4 μg/uL and 200 pmol/uL, respectively. For each nucleofection, we prepared complete Amaxa ‘P3 Primary Cell Solution’ (Lonza V4XP-3024) by adding 20 μL of the Supplement provided to 90 μL P3 Solution in a 1 mL sterile cryovial. Cas9 RNP was pre-assembled by mixing 2 μL of each sgRNA (16 μg, dissolved in T.E. buffer) and 2 μL of recombinant Cas9 protein (20 μg HiFi Cas9 nuclease v3; IDT), in a 1.5 mL sterile microcentrifuge tube and incubated at RT for 30 – 45 min. After 25 minutes, 0.8 - 1 X 10^6^ cells were harvested and added to the complete Amaxa ‘P3 Primary Cell Solution’. Immediately before nucleofection, 1 uL (200 pmol) of Alt-R HDR modified ssODN was added to the pre-assembled Cas9 RNP. This complete Cas9 RNP and ssODN was added to complete Amaxa ‘P3 Primary Cell Solution.’ We used the Amaxa 4-X unit nucleofector unit with a single cuvette (Lonza; V4XP-3024) filled with 100 μL of cells, ‘P3 Primary Cell Solution,’ and complete Cas9 RNP with bridge oligos (BOs). BO sequences are located in Methods, Table 1. The nucleofector program used was ‘Primary Cell P3′ program set using the pulse code ‘CG-104′. The nucleofected cells were washed with room temperature complete media containing HDR Enhancer V2 at 1 μM (IDT). The cells were cultured in a gelatinized 6-well plate at 32°C with 5% CO2 incubator for 1 day after which the media was replaced with complete meda lacking HDR Enhancer V2 daily, until the cells reached 60–80% confluency. During the first passage, genomic DNA was extracted from 1 X 10^6^ cells and we performed genotyping PCR with primers flanking the SV. Sanger sequencing was performed to confirm deletion. The remaining cells were diluted to 1 cell per 100 uL concentrations and pipetted into gelatinized 96-wells. Once distinct colonies formed from a single cell, the colony was passaged into a 12-wells plate. After reaching confluency, DNA and RNA were harvested for genotyping and assessing transcriptional effects.

### Sanger sequencing

PCR-amplified DNA was purified using a PCR gel purification kit (QIAGEN) and submitted to Quintarabio for Sanger sequencing with SV genotyping primers.

### Chromatin immunoprecipitation (ChIP) and ChIP-qPCR

ESCs (5-15 million) were fixed with 1-2% formaldehyde (Sigma-Millipore F1635-25ML) for 20 minutes at RT and quenched with 0.2 M glycine (Sigma-Millipore 50046-250G). The cells were pelleted at 288 x G for 5 minutes at 4°C, the supernatant removed, and the cells snap frozen until use. Cells were dounce homogenized on ice with a pestle, centrifuged at 845 G for 5 minutes at 4°C, and the supernatant was replaced with micrococcal nuclease (MNase) digestion buffer. After MNase digestion with Protease inhibitor and MNase (1500 units per 5 million cells), and sonication (200-260 uL of sample per tube using Diagenode Bioruptor (Diagenode) for 30”-on, 30”-off), ChIP was performed with 2-10 ug of DNA and 2 μg of antibody, followed by enrichment with protein G-Dynabeads (ThermoFisher Scientific 10004D) at 4°C for 3 hours. After stringent washing with High salt buffer, Tris/LiCl buffer, and finally Tris-EDTA buffer, DNA was eluted off the G-Dynabeads using Elution buffer. Proteins were degraded with Proteinase K (20 mg/mL, Ambion AM2546) at 65°C overnight. Samples were then purified with a Zymo PCR purification kit (Zymo Research D4014). Enrichment was measured by qPCR relative to 1% of input DNA. Primer sequences and buffer recipes are listed in **Methods, Table 1** and **2**.

### ChIP Library preparation and sequencing

ChIP DNA libraries were prepared with TruSeq ChIP Library preparation kit Set A (Illumina IP-202-1012) or ThruPLEX® DNA-Seq Kit (Takara R400675) and multiplexed using Nextera XT Index Kit v2 Set A (Illumina FC-131-2001) or DNA Unique Dual Index Kit Set A (Takara R400665). ChIP sequencing was performed on an Illumina Novaseq S4 to a depth of ∼50M reads per sample.

### ChIP Analysis

ChIP-seq data was analyzed with the ENCODE *chip-seq-pipeline2* v1.3.5.1. Reads were trimmed (Trimmomatic v0.39), aligned (Bowtie v2.2.6), and deduplicated (Picard - version 1.126). Peaks were called using SPP v1.14 (for transcriptions factors) or MACS v2 (for histone marks) and results were visualized in the Integrative Genome Viewer (Robinson et al., 2011).

### RNAseq and isoform quantification

Total RNA, with RNA Integrity (RIN) values > 8.5, A260/280 ratio ∼2.0 and A260/230 ratio ≥2.0, were processed into libraries using the KAPA mRNA HyperPrep Kit (Roche), fragmented to 250-300 bases, reverse transcribed, and ligated to Illumina sequencing adaptors prior to 10 cycles of PCR amplification. Libraries were sequenced on the Illumina Novaseq X Plus platform (150 bp paired-end reads, ∼100 million (100 M) reads per sample). The paired-end FASTQ files were aligned to the reference mouse genome (mm10) (Dobin et al., 2013) and quantified with RSEM (Li and Dewey, 2011) to obtain transcripts per million (TPM) .tsv files and outputs counts per million (CPM) values for all genes and transcript isoforms.

### Differential gene expression analysis

Gene counts were analyzed using DESeq2 R package (Love et al., 2014; R Core Team, 2023). Genes with adjusted *p*-values of < 0.05 and |log₂FC| ≥ 0.5 were considered differentially expressed.

### Differential exon usage analysis

Total RNA sequencing data was aligned to the *Mus musculus* GRCm38.86 genome build using STAR (Dobin et al., 2013). After generating count files, differential exon usage was quantified using the R package DEXSeq in Rv4.2.1 (Anders et al., 2012; R Core Team, 2023), which bins exons based on transcript annotations. The number of bins per genes is determined based on all transcripts from the mouse transcriptome, where exons of different length are separated into bins whose sum is the largest continual exon. Events with p-value ≤ 0.1 were considered significant.

### Exon usage qPCR validation

To validate exon inclusion, PCR was performed on cDNA using primers flanking constitutive and variable exons. Gel band intensities were quantified in ImageJ to calculate the percent spliced-in (PSI) ratio which is the intensity of the spliced exon band, divided by the intensity of a band corresponding to a stable isoform. Primer sequences are listed in **Methods, Table 1**.

### Data availability

CTCF ChIA-PET data for 129S1/SvImJ and C57BL/6J mouse ESCs were downloaded from the ENCODE portal (Sloan et al. 2016) (https://www.encodeproject.org/) with the following identifiers: ENCSR780SBE and ENCSR179IQN.

## ACKNOWLEDGEMENTS

We would like to thank Laura Reinholdt for providing the mouse embryonic stem cell lines. This study was funded by the Jackson Laboratory.

## AUTHOR CONTRIBUTIONS

S.S. and C.L. conceived the study. N.M. and S.S performed the mouse ESC experiments and analysis. M.K. provided cell pellets for ChIP experiments. S.S., C.L., N.M., and M.K. drafted and critically revised the article.

## Declaration of Interests

Charles Lee is a scientific advisor for Nabsys.

Sabriya Syed declares no competing interests.

Nicholas Moskwa declares no competing interests.

Minji Kim declares no competing interests.

## TABLES

Method Tables 1. Primers for SV genotyping, ChIP-qPCR, and gene expression

Method Tables 2. ChIP buffer recipes

Supplemental Table 1. SV information for 1,492 intronic CBLoop anchors

Supplemental Table 2. DESeq tables (B6 vs 129S)

Supplemental Table 3. DEXseq tables (B6 vs 129S)

---

**Supplemental Figure 1.**
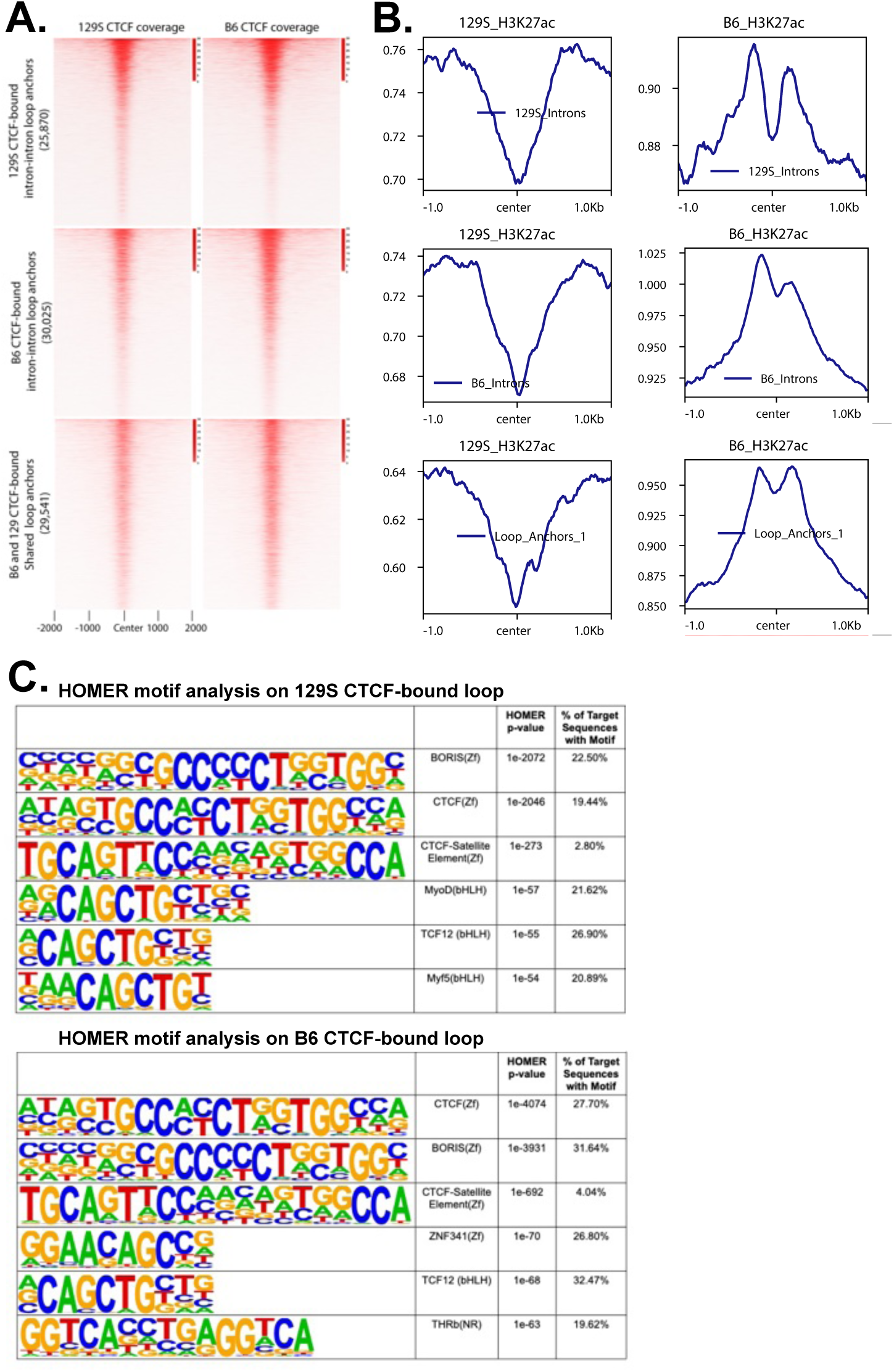
Characterization of a subset of intronic CTCF-bound B6 and 129S chromatin loops. **(A)** CTCF tag density plots displaying CTCF occupancy for CTCF-bound 129-specific intron-intron loop anchors (25,870), CTCF-bound intron-intron B6-specific loops (30,025), and loop anchors shared between the two strains (29,541); **(B)** Corresponding H3K27ac tag density plots for B6 and 129S around the intron-intron loop anchors displayed in panel A. Plot shows that H3K27ac flanks CTCF loop anchors; **(C)** Homer motif analysis for the intron-intron strain-specific CTCF loop anchors. The top 6 enriched motifs are displayed (target motif vs. background).

**Supplemental Figure 2.**
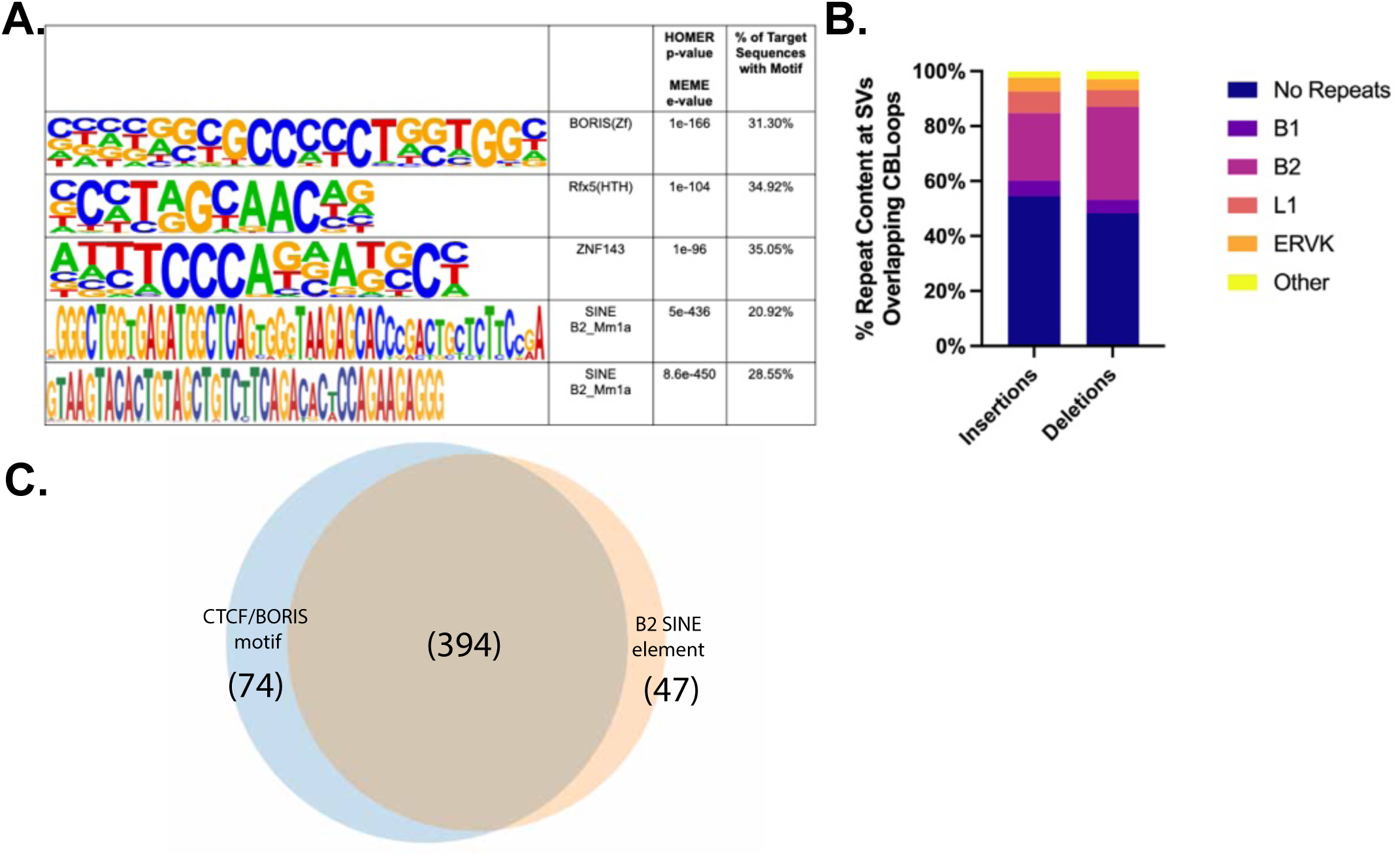
Repeat elements in intronic CTCF-bound SVs. **(A)** Homer and MEME motif analyses of the 1,492 SVs shows high incidence of B2 SINE elements; **(B)** Percent repeat content calculated for the SVs; **(C)** B2 SINE element containing SVs show overlap with the CTCF/BORIS motif containing SVs.

**Supplemental Figure 3.**
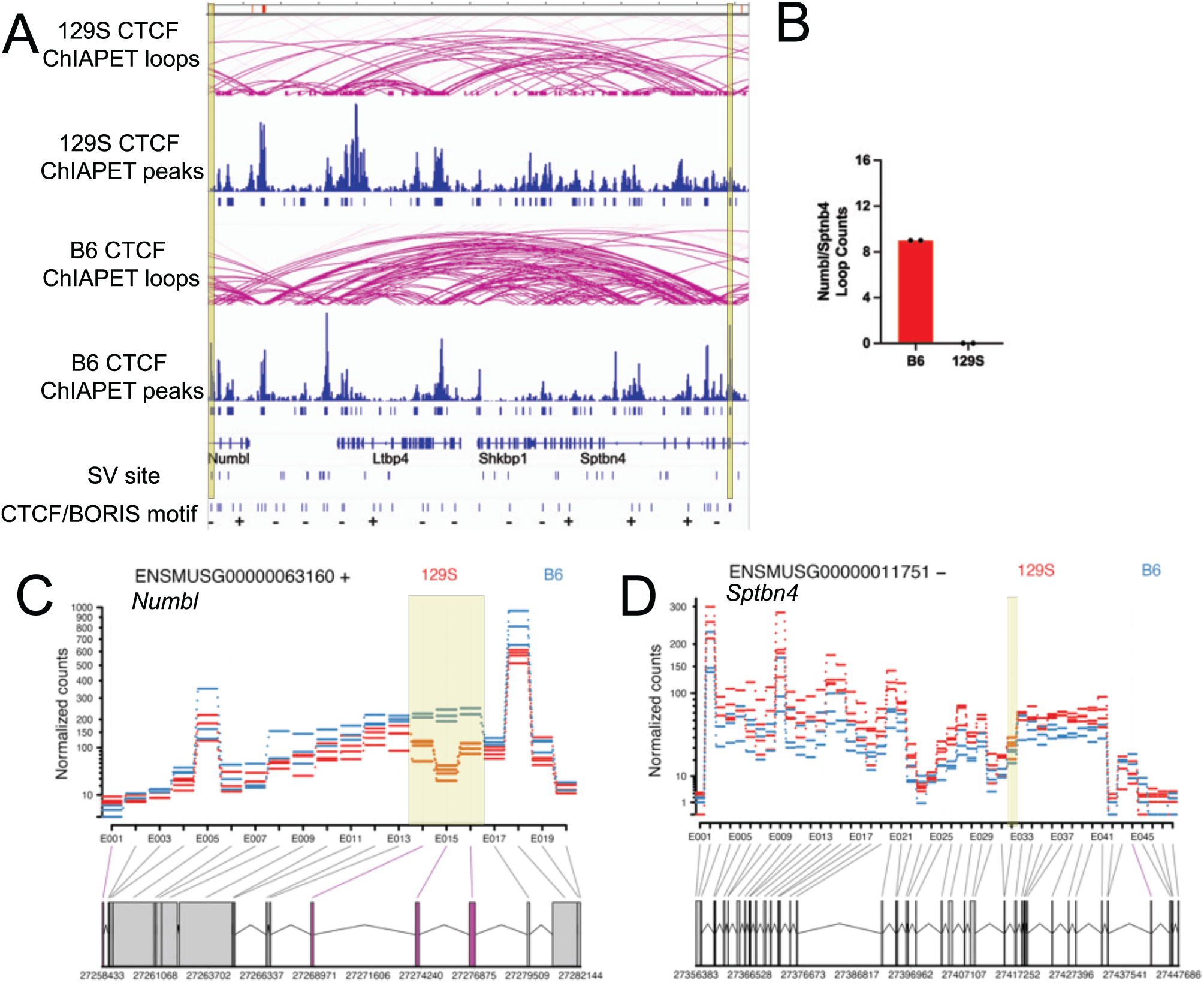
Differential exon usage for exons flanking SV chr7-26971377-DEL-261. **(A)** Genomic tracks (chr7:27,270,137-27,411,782) for loop bridging SV chr7-26971377-DEL-261 to gene Sptbn4 intronic region. Loop anchors highlighted in yellow; **(B)** Diffloop analysis shows the specific loop loss in the 129S ESC clones; **(C)** DEXseq normalized plot count showing significant differentially used exons in gene Numbl when comparing B6 and 129S ESC; **(D)** DEXseq normalized plot count showing significant differentially used exons in gene Sptbn4 when comparing B6 and 129S ESC.

**Supplemental Figure 4.**
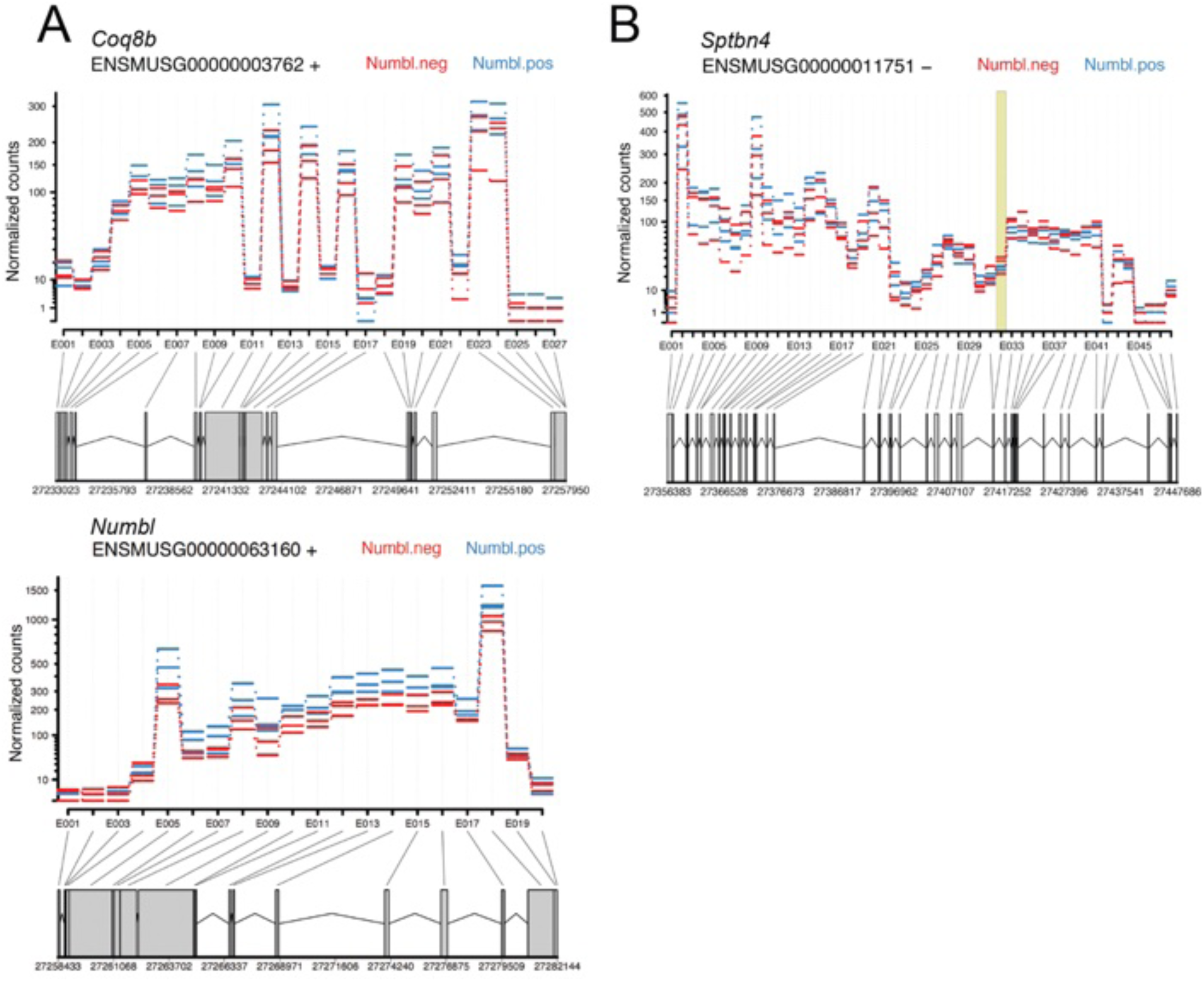
RNAseq of Numbl deletion clones validates differential exon usage observed in a B6/129S ESC comparison. **(A)** DEXseq normalized plot count showing no differentially used exons in gene Numbl and Coq8b, when comparing Numbl unedited to Numbl edited clones; **(B)** DEXseq normalized plot count showing differentially used exons in gene Sptbn4 when comparing Numbl unedited to Numbl edited clones.

**Supplemental Figure 5.**
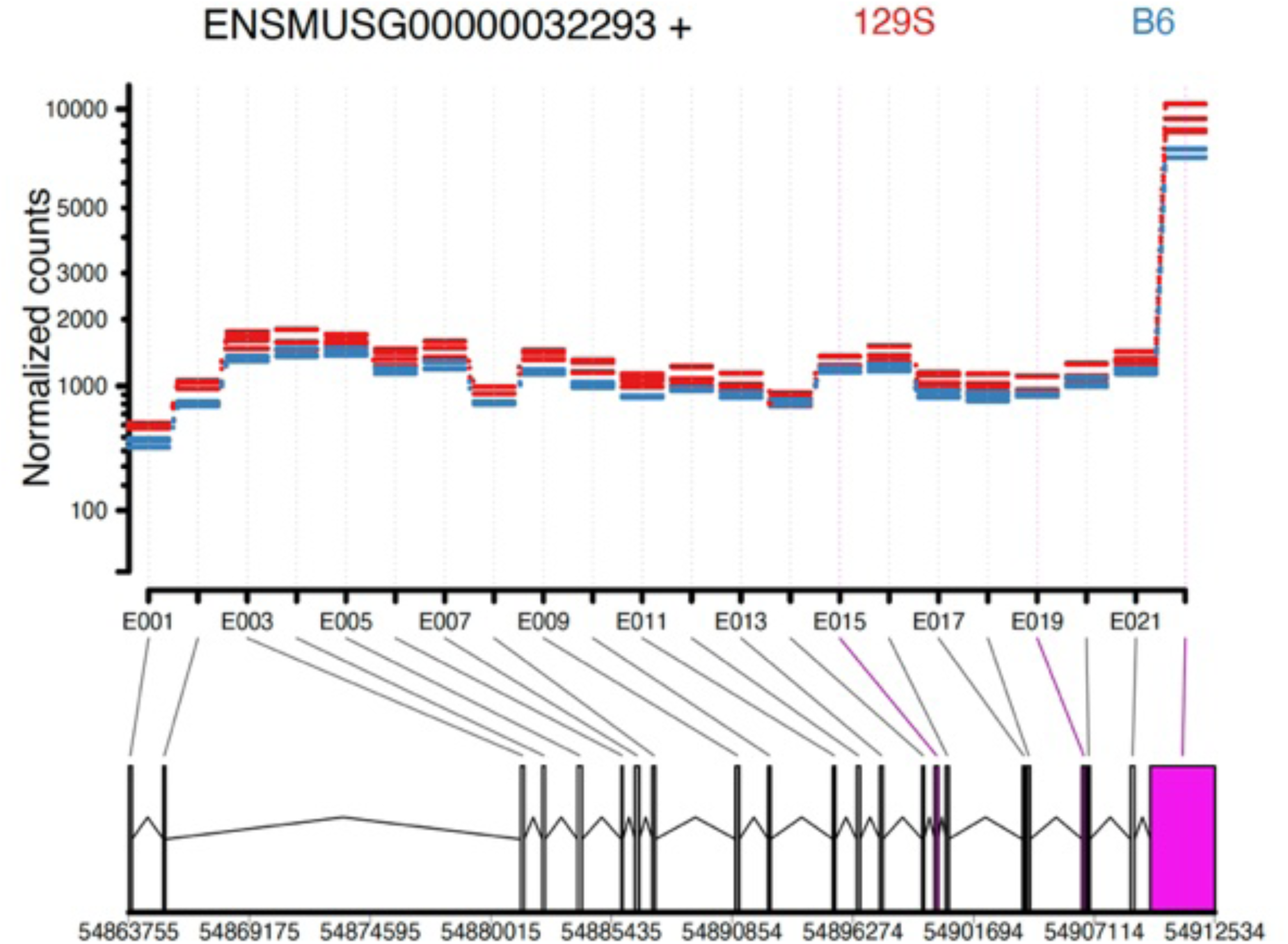
Differential exon usage for exons flanking SV chr9-54812235-DEL-668. DEXseq normalized plot count showing significant differentially used exons in gene Ireb2 when comparing B6 and 129S ESC.

## Notes

### Competing Interest Statement

Charles Lee is a scientific advisor at Nabsys. All other co-authors have no competing interests.

